# Novel method to determine diagnosis-defining refraction points

**DOI:** 10.1101/649442

**Authors:** Tsuneto Yamauchi, Mitsuhiro Ohshima, Yoko Yamaguchi, Kazunori Konishi, Kai Kappert, Shigeru Nakano

## Abstract

Diagnosis of a certain disease generally relies on definitions established by professional medical societies and comprise the patient’s history, physical examination, and test results. These include physical compositions such as body mass index (BMI), and laboratory tests such as serum creatinine and albumin in urine samples. In general, laboratory tests are based on mathematical methods, *e.g.* defining critical values from the mean ± kσ of a population, where k is a natural number and the standard deviation is σ (“mean ± kσ-method”). In most cases k is defined as 2, leading to reference ranges defining 95% of test results as normal. However, this method mostly depends on a normal distribution of values.

Here we applied a novel method (“SoFR-method”) based on data sorting to define refraction points, which carry informative value as diagnostic criteria. Applying the SoFR-method, standard measures such as critical BMI-values are categorized by equal robustness as by the mean ± kσ-method. However, the SoFR-method showed higher validity when analyzing non-normalized values such as creatinine and albumin, as well as hepatocyte growth factor (HGF) and hemoglobin in a novel Perioscreen assay in saliva of diabetic and non-diabetic patients.

Taken together, we defined a novel method based on data sorting of test results from patients to effectively define refraction points which might guide more accurately clinical diagnoses and define relevant thresholds.

## Introduction

The determination of a pathophysiological condition or the diagnosis of a certain disease usually rely on definitions established by professional medical societies. These diagnostic criteria are based on the combination of patient’s signs, symptoms, thus patient’s history, and physical examination and test results [1, 2]. Test results are quantifiable and comprise measures such as body weight and body mass index (BMI) as well as laboratory tests such as serum creatinine and albumin in urine samples.

Deducing diagnostic value from laboratory test results depends on mathematical methods [3, 4]. These define – presumably – critical values from the mean ± kσ of a healthy population, where k is a natural number and σ is the standard deviation. Depending on the rigorousness desired, k varies from 1 (epidemiological measures) to 3 (tumor markers and cardiac troponin), but often is 2 (most laboratory tests, such as transaminases, glucose, lactate dehydrogenase etc.), where the range within these thereby defined 95% is considered the reference range [5–7]. Indeed, the broadly used current practice defines this reference interval based on collected ~120 samples derived from healthy individuals in a population [3]. Despite uncertainties determining “healthy” in cohorts, values between the 2.5^th^ (almost equal to mean − 2σ) and 97.5^th^ percentiles (almost equal to mean + 2σ) or the 99^th^ percentile (mean + 3σ) are used for the definition of a reference interval.

Recent mathematical attempts were undertaken to more specifically address population-based issues, in particular in populations with high percentage of various races and ethnicities, such as in the United States [8, 9]. When using large data basis, such as electronic health records [10], reference intervals are sharpened and display differences between demographics, which might have clinical impact including prognostic value.

Certainly, the reference range including the upper and lower reference limit guides clinicians to interpret test results, knowing that these 95% of values are found in healthy individuals. Even though this method is broadly acceptable and accurate, in particular if the standard deviation is not very large compared to the mean, this method, however, largely depends on a normal distribution of values [7]. Thus, taken into account that *e.g.* laboratory tests display integral components for medical diagnoses and decisions in patients, the mean ± kσ method is limited in case of non-normal distribution of quantified clinical data. While large data sets might be useful in more accurately accounting for individual patients’ demographics [9, 10], they still rely on normal value distributions and follow rather strict mathematical paradigms, which are prone to overlook better and more clinical relevant refractions points displaying changes in physical conditions, and therefore are critical for individual diagnoses. Here we applied a novel method based on consecutive data sorting visualizing such refraction points. In particular, this novel method is applicable in case of non-normal distributed data and, in contrast to the mean ± kσ method, is based on data sets containing healthy and diseased individuals.

## Methods

### General demographics and routine examinations

Data from n=803 patients were obtained in the Shika-Clinic, Ishikawa-prefecture, Japan, between September 2017 and March 2018. All patients enrolled in this study documented informed consent. N=400 patients were males, and n=403 patients were females. The age ranged from 27 to 92 (mean 67.64±10.22) years. In a subset of analyses for periodontitis screening, n=307 patients were enrolled. Of these, n=121 of were diagnosed with type 2 diabetes and n=186 of included patients were non-diabetics.

All patients underwent routine clinical examinations, including determination of body weight and height. The body mass index (BMI) was calculated according to the following definition: BMI = body weight (kg) / body height (m)^2^.

This study was approved by the Ethics Committee of the Ohu University (#194), and Nihon University School of Dentistry (#EP17D007-1) and the study was conducted in accordance with the Helsinki Declaration of 1975, as revised in 2013, and the ethical guidelines for epidemiology research of 2002, as revised in 2008 (Ministry of Health, Labor and Welfare, Japan).

### SoFR-Method

Here, we define the method which will show the refraction points of the data. From the original data, we will sort it in ascending order. When displayed on a graph, it may be possible to find refracted points. This method for finding refraction points, we will call “SoFR-Method” (this is an abbreviation of “Sorting, Finding the Refraction Points”).

### Periodontitis screening

Screening for periodontitis (“Perioscreen” [11]) was done in n=648 patients in saliva samples. Based on combinatory analyses with HGF levels, a final cohort of n=174 data points were used for analyses. Each patient was instructed to rinse with 3 ml of water in the mouth for 10 s. Afterwards, the samples were collected in a paper cup. Saliva occult blood test was carried out using a test paper strip assay [11]. Briefly, an anti-human hemoglobin monoclonal antibody with colloidal gold conjugate was immobilized on membranes as capture antibody, followed by adding of saliva specimen. Based on capillary movement and membrane chromatography, positive reaction results from complex binding and subsequent red-violet color reaction. The test strip was shortly placed into 1:4 diluted saliva. The immunocomplex was visualized after 5 minutes. The sensitivity of the assay was determined at ≥ 2μg/ml. Results were classified as grades 0 to 5.

### Nephropathy screening

From n=482 patients urine was taken to evaluate potential prevalence of nephropathy. Based on combinatory analyses with HGF levels, a final cohort of n=132 data points were used for analyses. Urinary albumin in spot urin was measured by immunonephelometry using a kit from NITTOBO MEDICAL CO., LTD. (Tokyo, Japan).

Serum and urinary Cr was measured using enzymatic assays (SEKISUI MEDICAL. CO., LTD., Tokyo, Japan).

### Quantification of hepatocyte growth factor and hemoglobin in saliva

Besides periodontitis screening, collected saliva (mouth rinse) samples were used for quantification of hepatocyte growth factor (HGF) and hemoglobin in n=318 and n=322 patients, respectively. HGF concentration in rinse water samples was measured by using HGF ELISA kit (Quantikine human HGF immunoassay, R&D systems, Minneapolis, MN, USA) according to the manufacturer’s instructions.

The rinse water samples stored at −80 °C were thawed on ice and diluted (2.86 to 3 folds, depending on the residual volume of each sample) with distilled water. Hemoglobin (Hb) concentration in rinsed water was measured using a commercially available kit for latex agglutination turbidimetry (LZ test - Eiken HbAo, Eiken Chemical Co., Tokyo, Japan) according to the manufacturer’s instructions. In this system, Hb measurement in saliva should be performed after diluting saliva with a specific buffer solution which stabilizes the Hb before measurement. However, in the present study, the samples were originally diluted and the specific buffer solution was not employed.

### Statistics

All data points were sorted ascendingly by its magnitude. After sorting the data, the points where the trends of the sorted data were visually changed with regard to its direction, we will call these points “refraction points.”

On the other hand, the “Inflection point” is mathematically defined. This is defined as the zero points of the second derivative of a smooth curve. In the case of the normal distribution, there are two inflection points which are ± 1 x σ away from the mean (where σ is the standard deviation of the normal distribution).

Further, for testing the normality of data, Shapiro-Wilk normality test was applied. In the Shapiro-Wilk normality test, the null-hypothesis is the data are come from a normally distributed population. If the p-value is less than chosen level of significance, then the null-hypothesis is rejected. On the other hand, if the p-value is greater than the chosen level of significance, we can’t refuse the null-hypothesis. In the case of the level of significance 0.05, if p-value >0.05, then the null-hypothesis can’t be rejected.

In this paper, for numerical analyses and graph drawing, the statistical software “R” was used.

### Results

In this part, we explain four examples of using the SoFR method. The Shika-Clinic data was collected for the research of the relationship between dental health and diabetes. We study two cohorts; diabetic patients and non-diabetic patients. We study these two cohorts in terms of the HGF and Hemoglobin in saliva.

We are also searching the reference intervals of the values of the HGF and Hemoglobin in saliva. It is very hard to find the thresholds, but the first two examples show that SoFR method will suggest several candidates of the thresholds.

The last two examples are the simple applications of the SoFR method to albumin-to-creatinine ratio (ACR) and BMI (Body Mass Index).

### Hepatocyte Growth Factor in saliva

Using the Perioscreen assay [11], we analyzed both hemoglobin and HGF concentration in saliva collected in patients, separated into cohorts of diabetics and non-diabetics. As shown in Figure 1A, HGF was determined within a range of 26.4 pg/ml and 13574.1 pg/ml, with large variation and significant differences between the 2 cohorts. (Fig 1A)

**Figure 1A.**
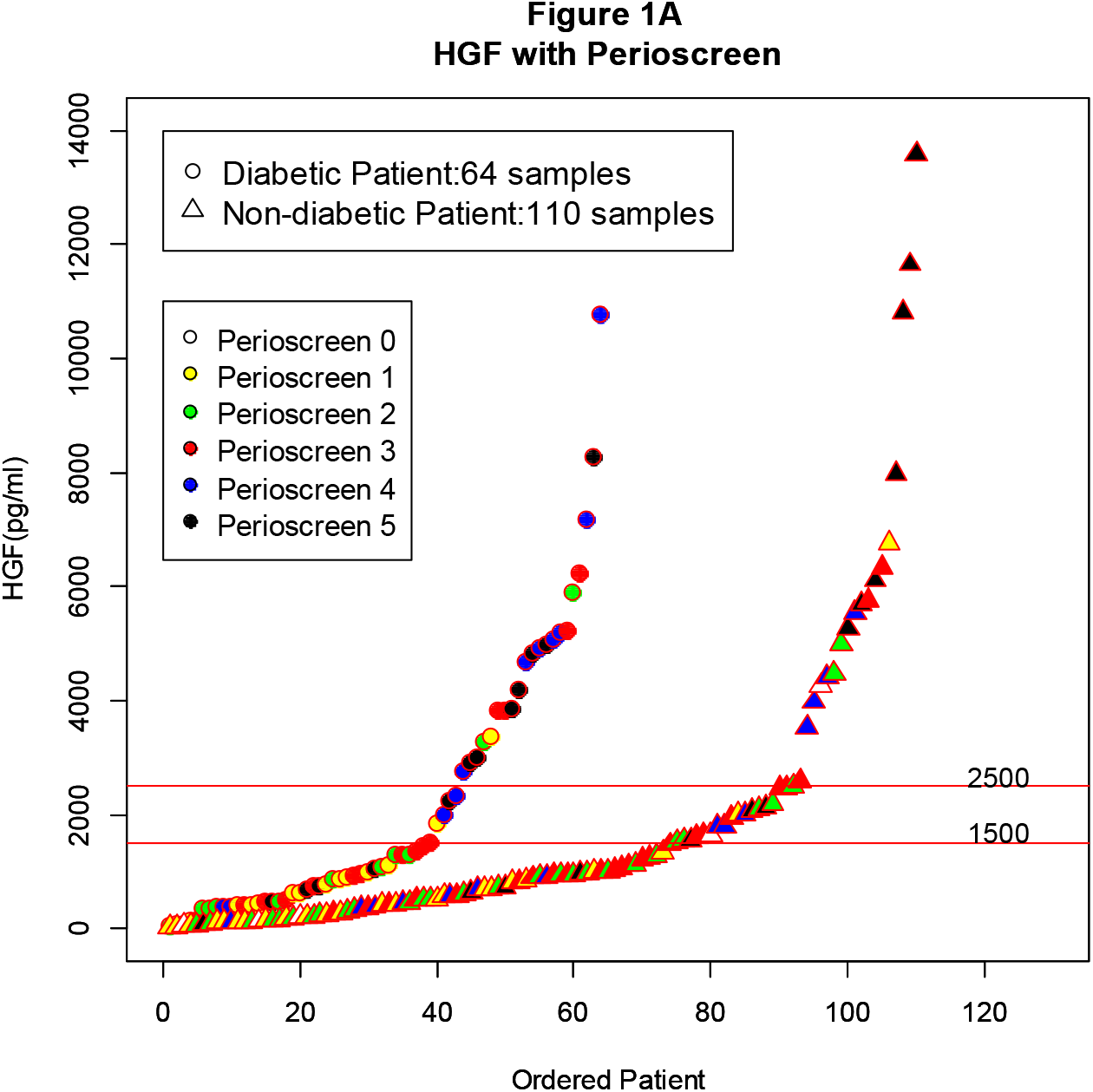
HGF with Perioscreen. It is the graph which showed the measurement of HGF and the relation of the value of a perioscreen. When the HGF measurement values are sorted in ascending order, divided into diabetic patients and non-diabetic patients, it is visually shown that the value of Perioscreen remarkably increases as the HGF value increases. Furthermore, it can be seen that the location of the refraction points of HGF differ between diabetic and non-diabetic patients, which is another new question.

While in non-diabetic patients the visualized refraction point was determined at 2500 pg/ml, diabetics were characterized by a refraction point of 1500 pg/ml. Next we dichotomized each cohort into two subsets of patients:

HGF levels < 1500 pg/ml (subset A) and HGF levels ≥ 1500 pg/ml (subset B),
HGF levels < 2500 pg/ml (subset C) and HGF levels ≥ 2500 pg/ml (subset D) (Table 1).

**Table 1:** HGF values and Perioscreen.

| | Perioscreen<br>level $\leq 2$ | Perioscreen<br>level $\geq 3$ | Perioscreen<br>level $\leq 2$ | Perioscreen<br>level $\geq 3$ |
| --- | --- | --- | --- | --- |
| Subset A (HGF < 1500) | **21/38** | **17/38** | **55.3%** | **44.7%** |
| Subset B (HGF ≥ 1500) | **4/26** | **22/26** | **15.4%** | **84.6%** |

**Non-diabetics**
| | Perioscreen<br>level $\leq 2$ | Perioscreen<br>level $\geq 3$ | Perioscreen<br>level $\leq 2$ | Perioscreen<br>level $\geq 3$ |
| --- | --- | --- | --- | --- |
| Subset A (HGF < 2500) | **54/92** | **38/92** | **58.7%** | **41.3%** |
| Subset B (HGF ≥ 2500) | **4/18** | **14/18** | **22.2%** | **77.8%** |

As for the HGF example, we introduced another medical characteristic, which is the Perioscreen level at the following definition: Perioscreen ≤ 2 representing good oral condition and Perioscreen ≥ 3 defining bad oral condition. In diabetics only 15.4% were characterized at good oral condition with simultaneously HGF levels ≥ 1500 pg/ml, while the vast majority with enhanced HGF levels displayed a Perioscreen level ≥ 3. Strikingly, almost the equivalent proportion of a Perioscreen value, separated at ≤ 2 and ≥ 3, was seen in non-diabetics when their cohort-specific refraction point of 2500 pg/ml of HGF was applied.

As shown in Figure 1B, the QQ-plots demonstrate lack of normal distribution of this data set, which refuses the necessity to apply distribution functions or applying the mean ± k x σ method. (Fig 1B-1 and 1B-2)

**Figure 1B-1 and 1B-2.**
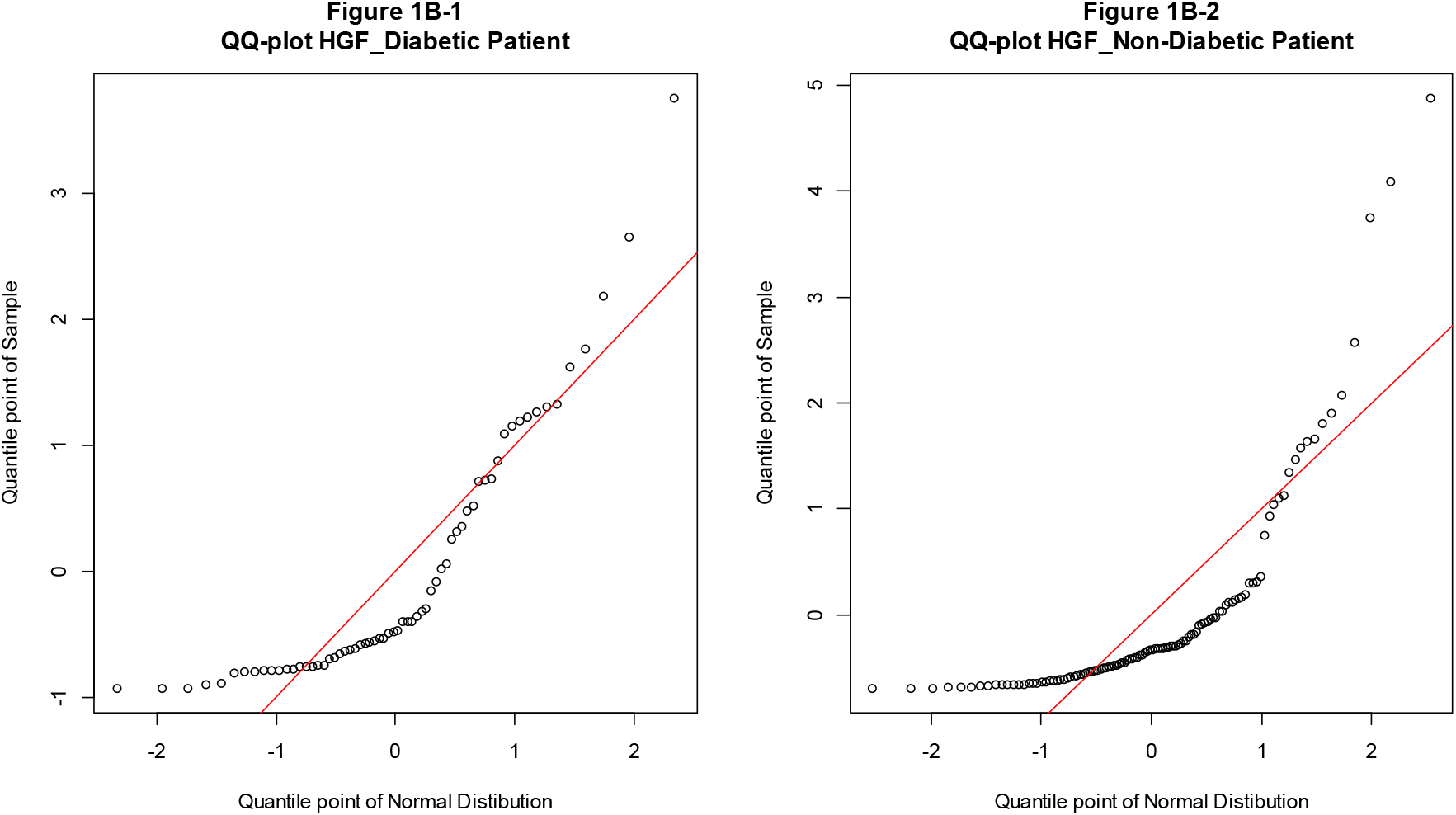
QQ plots HGF. QQ plots were generated to check whether HGF values follow normal distribution for diabetic and non-diabetic patients. In any case, normality is denied, and visual judgment is considered to be superior to forcibly creating a reference interval with an mean ± kσ.

### Hemoglobin in saliva

We further examined hemoglobin concentration in saliva specimen.

Following, we sorted the data and colored values from the Perioscreen assay are shown in Figure 2B. We detected different refraction points for both analyzed cohorts. (Fig 2A) While we observed a refraction point at 2.0 μg/ml for diabetics, there were two refraction points at higher values for non-diabetic patients, at 4.0 μg/ml and 7.0 μg/ml, which might reflect distinct pathophysiological changes for this study group. Therefore, we dichotomized each cohort into two subsets of patients:

hemoglobin values < 2.0 μg/ml (subset A) and ≥ 2.0 μg/ml (subset B) for diabetics,
hemoglobin values < 4.0 μg/ml (subset C) and ≥ 4.0 μg/ml (subset D) for non-diabetics.

**Figure 2A.**
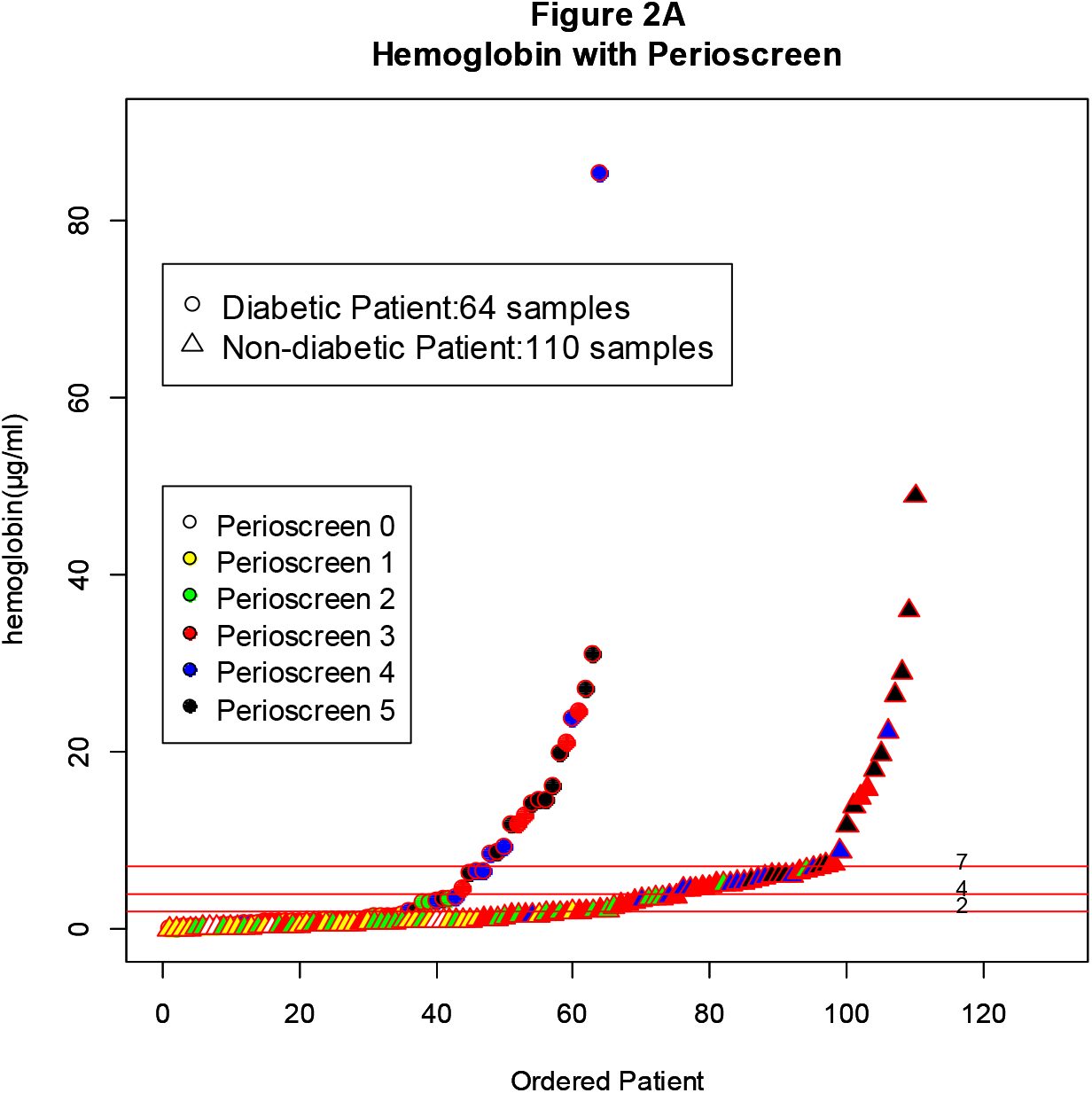
hemoglobin with Perioscreen. The same analysis as HGF was carried out for the relationship between hemoglobin value and perioscreen. Again, the refraction points differ between diabetic and non-diabetic patients. Moreover, it can be visually observed that the value of the Perioscreen increases as the hemoglobin value increases.

**Figure 2B-1 and 2B-2.**
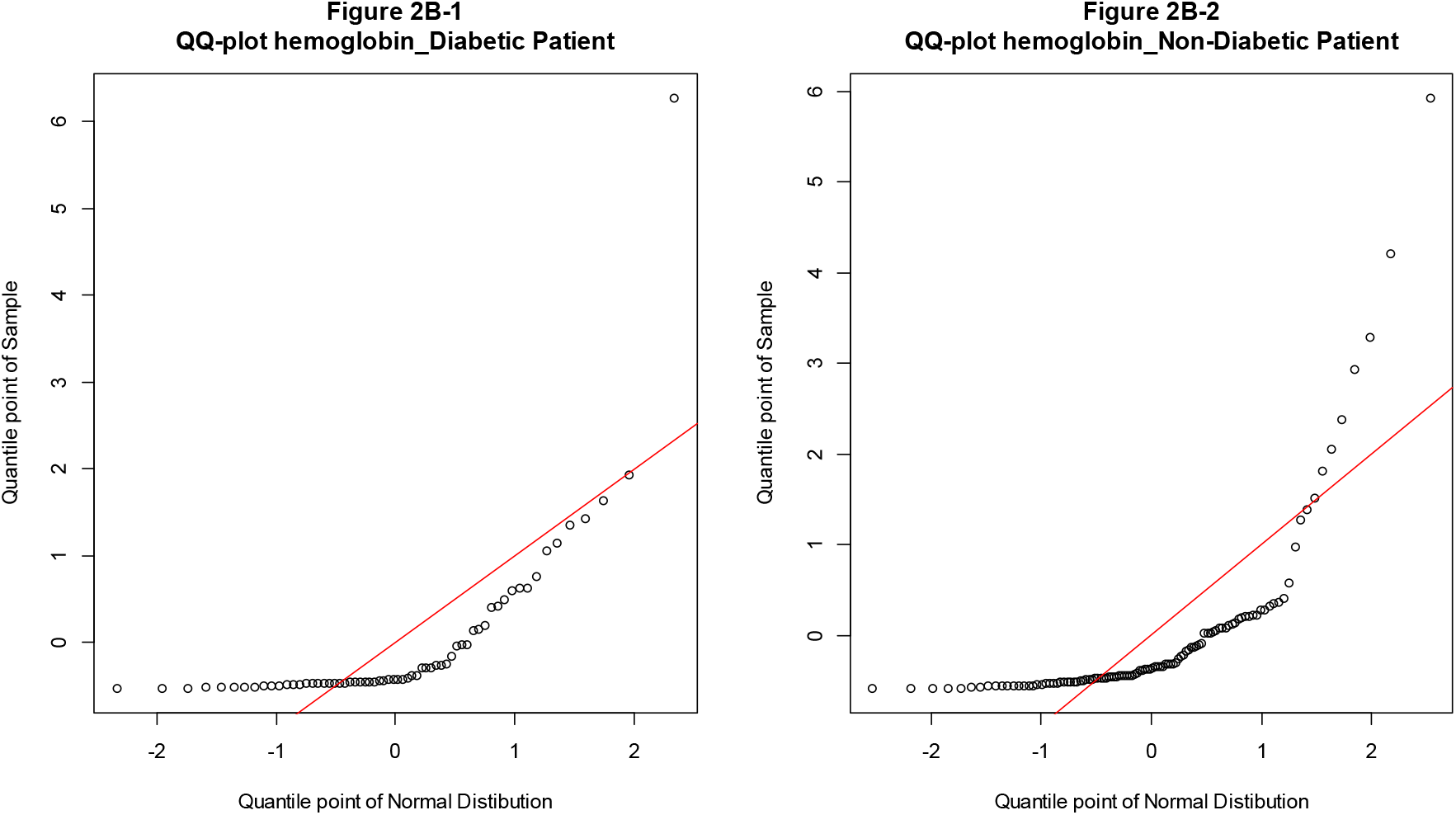
QQ plots hemoglobin. QQ plots were generated to check whether hemoglobin values follow normal distribution for diabetic and non-diabetic patients. In any case, normality is denied, and visual judgment is considered to be superior to forcibly creating a reference interval with an mean ± kσ.

As for the HGF analyses, we introduced the Perioscreen level as another medical characteristic, following the definition Perioscreen ≤ 2 representing good oral condition and Perioscreen ≥ 3 defining bad oral condition. Table 2 shows that among diabetics the vast majority with hemoglobin values above 2.0 μg/ml displayed Perioscreen levels ≥ 3 (88.9%) and, thus, were characterized by bad oral conditions. Non-diabetics, when separated into subsets at the significantly lower hemoglobin value of 4.0 μg/ml, also the highest proportion of patients displaying a Perioscreen level ≥ 3 were found in the group of higher hemoglobin concentration in saliva (94.3%). (Table 2)

**Table 2:**
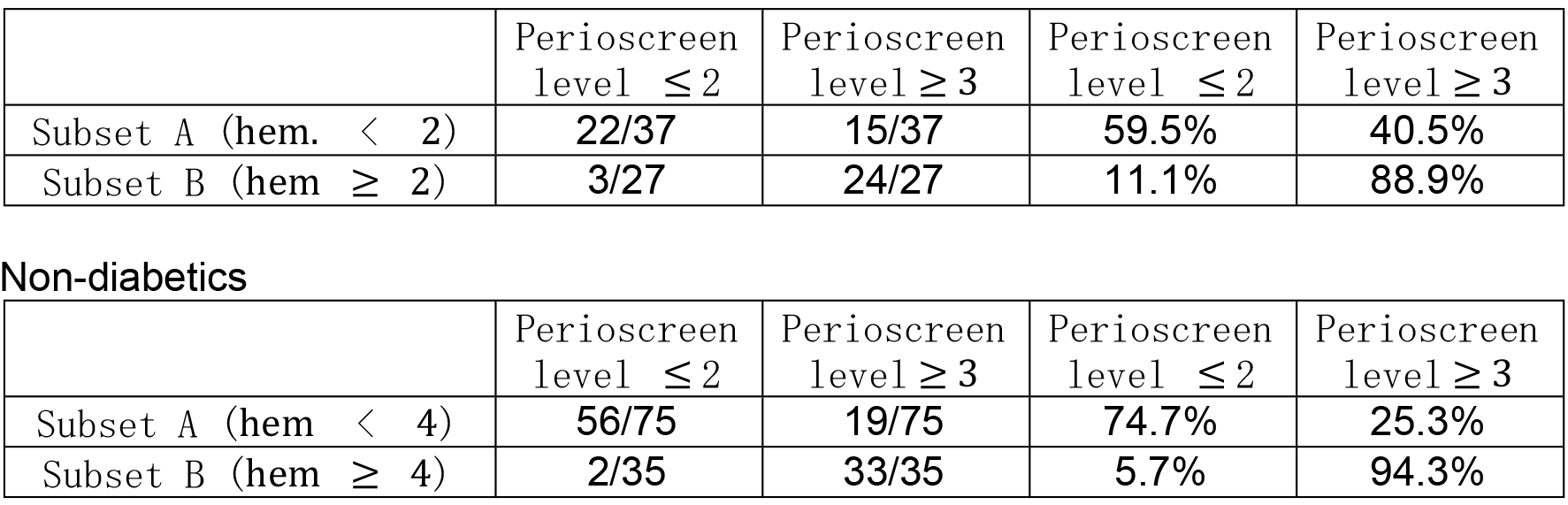
hemoglobin values and Perioscreen.

As shown in Figure 2B, the QQ-plots demonstrate lack of normal distribution of this data set, which refuses the necessity to apply distribution functions or applying the mean ± k x σ method. (Fig 2B-1 and 2B-2)

### Albumin-to-creatinine ratio in urine

Next we applied the SoFR in another data of the Shika-Clinic, including evaluation of urine albumin-to-creatinine ratio (ACR), well-established to detect early kidney disease in diabetic patients or patients at high risk for development of renal insufficiency like hypertensives [13].

The original data contains samples with values of 1556.60. It is not an outlier, but we would like to ignore this sample for research purposes. As shown in Figure 3A, a refraction point of ACR was visualized at 30mg/g creatinine for n=123 samples when data were evaluated where HGF-screening was performed. (Fig 3A)

**Figure 3A.**
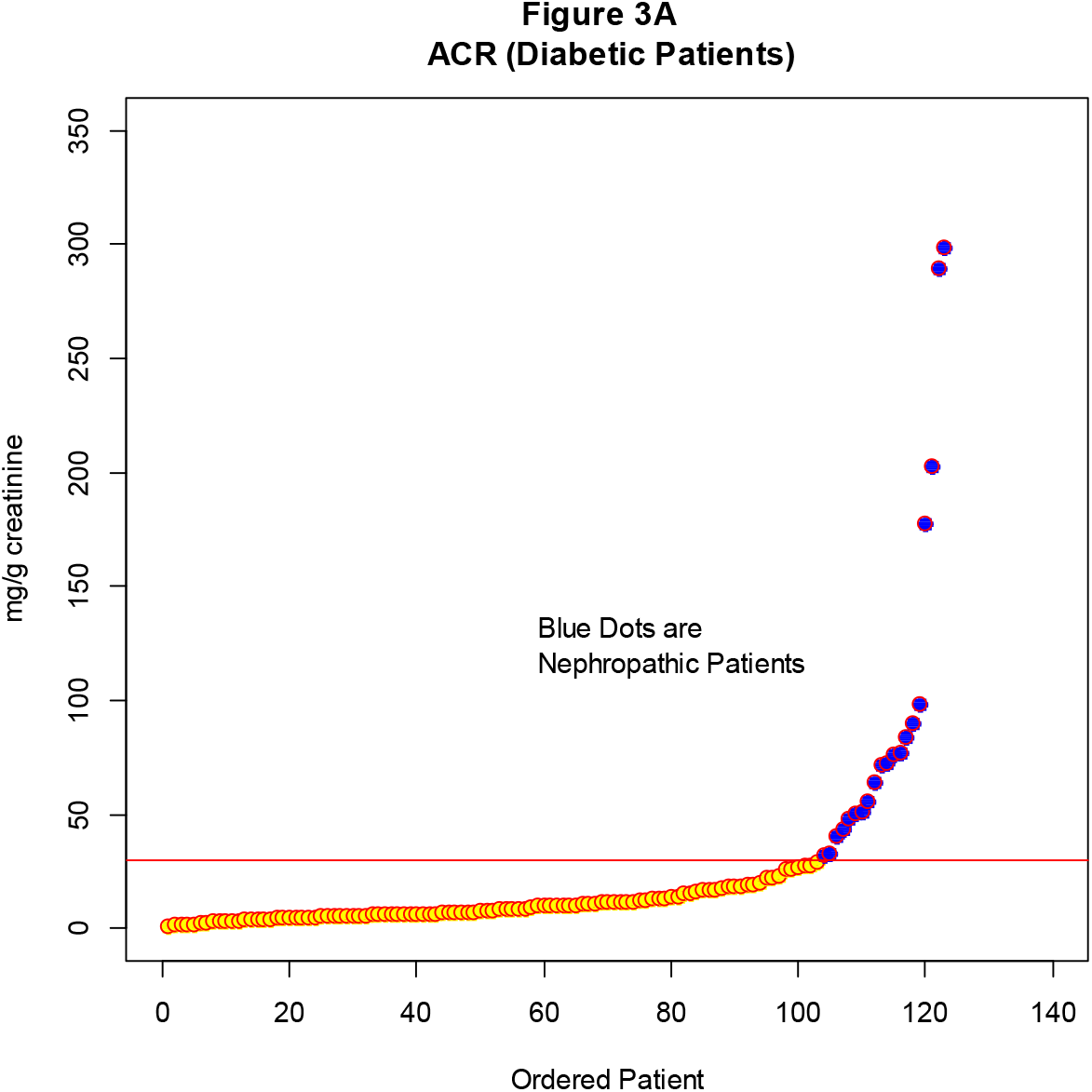
ACR (Diabetic Patients) It is the graph which arranged the ACR of a diabetic patient in order of size, and looked at the refraction point. There is a change point around 30 mg/ml, at this point it is possible to distinguish whether or not the patient is nephropathy even in actual diagnosis.

We further separated the data at a refraction point α with “A” being the subset of data with values <α and “B” as the complementary data set of “A“ with values > α. In the case of ACR, α is 30.0 mg/g creatinine with “A” as the subset of patients characterized by ACR-values <α (< 30.0 mg/g creatinine). “B” is the subset of patients with ACR-values ≥ 30.0 mg/g creatinine. By introducing another medical characteristic, such as nephropathy, we show that the number of patients with nephropathy in “B” is much greater than in “A”, underlining the usefulness of the SoFR-method.

In contrast, applying the “mean ± σ method”, the mean of creatinine data of the Shika-Clinic was calculated to 24.32 mg/g creatinine, 1σ to 45.76 mg/g creatinine and thus ”mean + 1σ” equaled 70.08 mg/g creatinine. Moreover, for the minus part of “mean ± σ method”, we will get minus value (−21.44), it is not acceptable. As visualized in Figure 2A, there is no evidence for a refraction point at the mean + 1σ value, suggesting inferiority of the “mean ± 1σ method” compared to the SoFR-method.

Figure 2B shows the QQ-plot of the ACR data. Apparently, ACR data are not normally distributed. (Fig 3B)

**Figure 3B.**
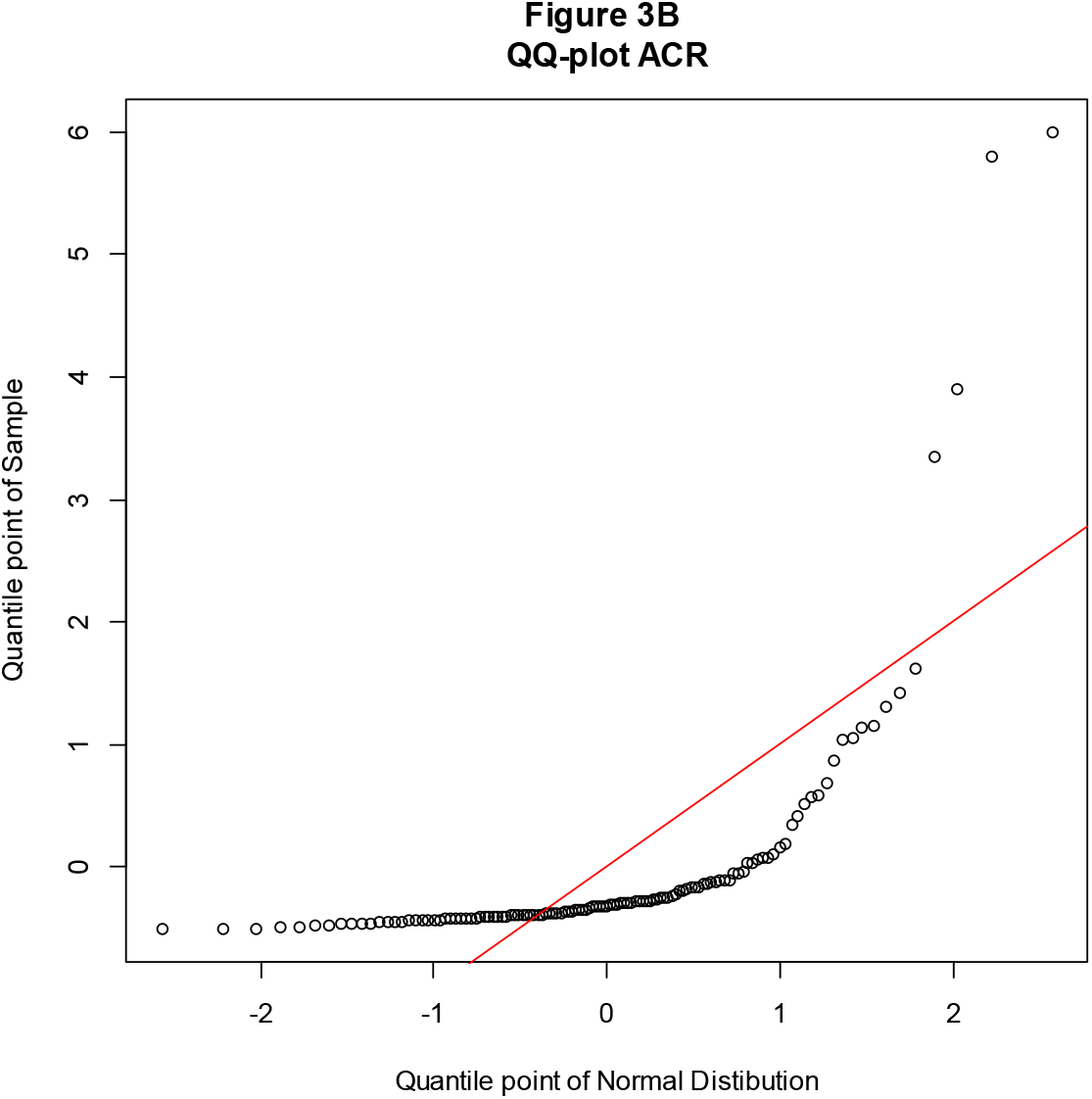
QQ-plot ACR. It was checked by QQ plot whether ACR value is normally distributed. It can not be said that it is a normal distribution.

### Body weight and Body Mass Index

Body weight was measured in all 694 patients and the BMI was calculated. Lowest BMI was 15.5 and highest BMI was 43.2. Thereafter, data was sorted by the BMI values in ascending order, depicted in Figure 4A, showing sigmoidal shape of the data and two refraction points at 20 and 30. In Figure 4A also the calculated mean ± 1 x σ (red lines) and BMI=20 and 30 (blue lines) are visualized. According to classifications by the WHO normal weight considered in the range of 18.5 to < 25 and obesity at values ≥ 30 [12], thus at the same value as we determined the second refraction point after data sorting. (Fig 4A)

**Figure 4A.**
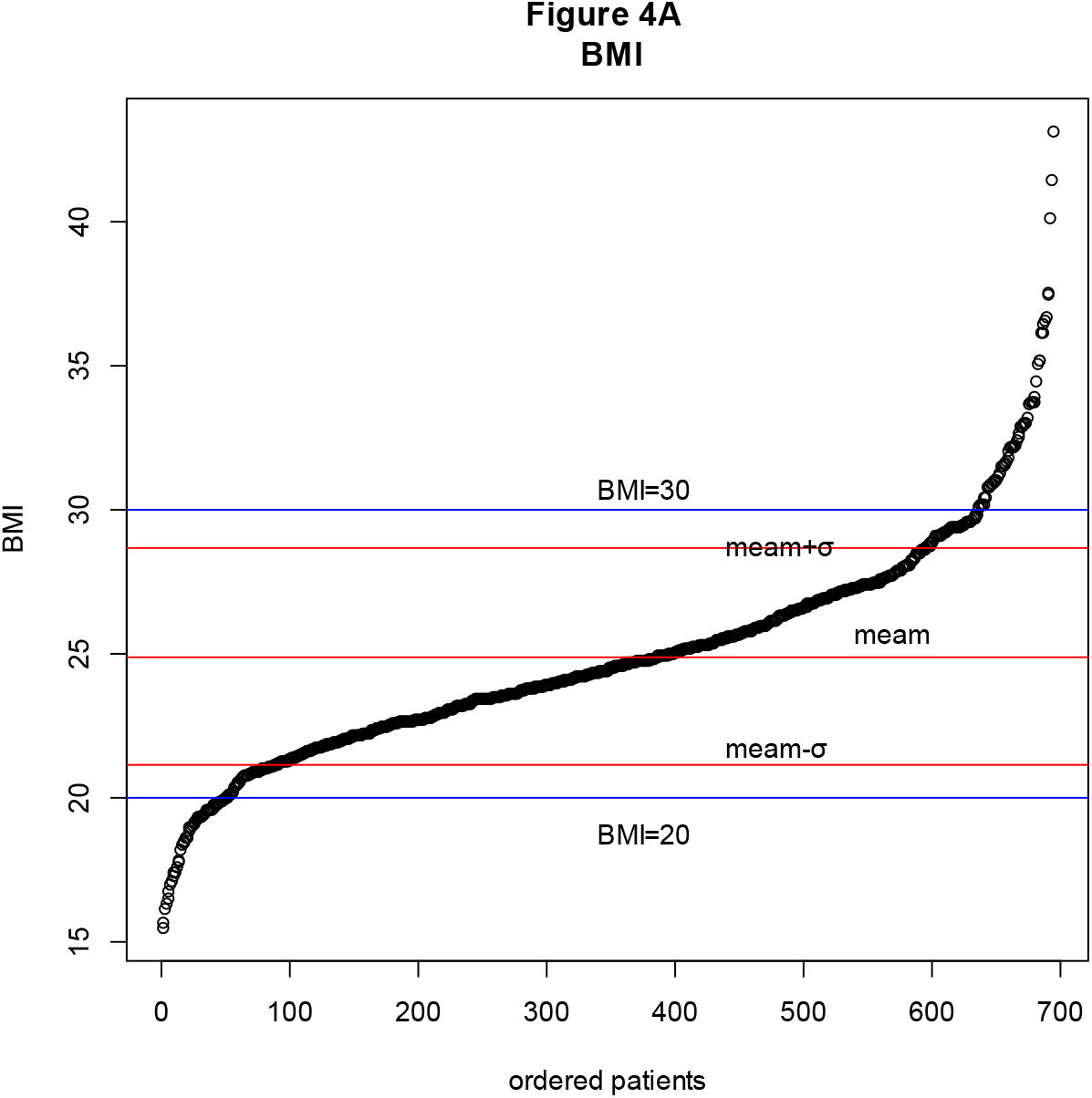
BMI. The Shika-Clinic data contains 694 patients’ BMI data. This is a graph in which BMI data are arranged in order of size and the refraction points are seen.

We next performed Quantil-Quantil (QQ) plots of BMI values for quantile points of normal distribution and quantile point of samples (Figure 4B). The graph visualizes data distribution and shows the degree of the normality of BMI data. However, it is difficult to justify the normality from this graph. (Fig 4B)

**Figure 4B.**
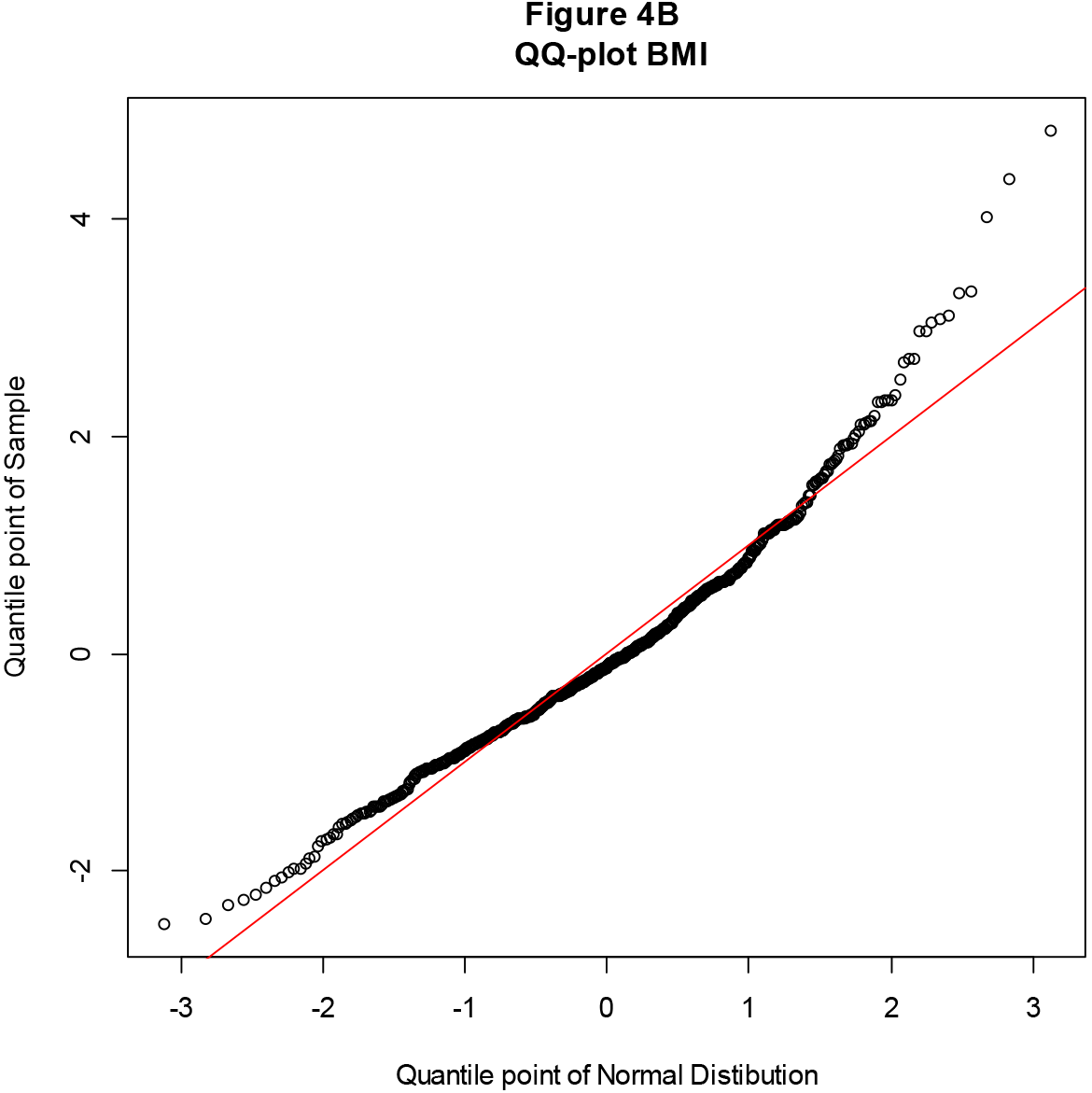
QQ-plot of BMI. QQ plots were generated to ensure that the BMI data followed the normal distribution. According to this, the central part shows a good fit to the normal distribution, but not the fit at both ends. Therefore, Shapiro-Wilk normality test was performed, the result is that normality is denied.

Using the Shapiro-Wilk normality test the p-value of 4.18 x 10^−12^ < 0.05 was calculated. Thus, this test shows that the BMI data does not follow a normal distribution. Thus, SoFR, which is based on visual recognition of refraction points, is superior to the argument assuming the normal distribution.

## Discussion

Here we demonstrate that the SoFR-method, based on plotting data in study groups or cohorts, is useful to define refraction points, which can be used for thresholds in clinical settings. We show applicability the SoFR-method in case of non-normal distributed data for 4 different conditions: BMI, ACR, and both hemoglobin and HGF in saliva. Thus, the SoFR-method might be useful for professional clinical assessment or referring third data, evaluating the patient’s physical condition.

While diagnosis of pathophysiological conditions conventionally relies on established definitions, data of quantifiable conditions are not always normally distributed among cohorts or study groups [14, 15]. Since the mean ± kσ method shows limitations in case of non-normal distribution of quantified data sets, here we established a novel method to define refraction points, visualizing changes in clinical conditions. This SoFR-method is potentially useful to establish cut-off values beyond the established mean ± kσ method.

Sorting BMI data in ascending order and visualizing refraction points from a cohort of 694 patients enrolled in this study, we observed a sigmoidal shape of the data with refraction points at 20 and 30 kg/m^2^. Importantly, also the WHO defines obesity at 30 kg/m^2^ for adults [16], which underlines the validity our sorted data. However, it has to noted that in contrast to *e.g.* clinical-chemistry data from blood samples, the WHO definition is not extracted from the normal healthy population, but was rather defined in terms of recommended values, easily to be communicated from clinicians to their patients [12].

As for BMI, ACR values were also not normally distributed among our cohort. Nonetheless, by sorting the data in ascending order we detected a refraction point at ACR30, which, strikingly, represents also the cut-off used in clinical diagnostic practice [17, 18]. This clearly shows that our SoFR-method is not susceptible to mathematical conditions such as normal-distribution, in contrast to the mean ± kσ method.

Further, here we analyzed the clinical data of a novel assay to screen for periodontitis disease: Perioscreen [11, 19]. We determined hemoglobin and HGF as parameters in saliva and sorted our data with regard to Perioscreen values in patients divided into diabetics and non-diabetics based on the significance of HGF for development of periodontitis [20, 21]. Regarding HGF, the SoFR-method strongly argues for the usefulness of defining refraction points by plotting and visualizing data, since we observed refraction points, which differed in magnitude between diabetics and non-diabetics. Moreover, since we introduced the clinical entity of oral condition (Perioscreen value) we observed differences in refraction points between the study groups which could be taken into account for applying cut-off values for individual patients. This suggests that susceptibility to periodontitis in diabetic patients might be triggered by HGF and also depends on differences in HGF values. Our data on HGF were supported by the finding that also significant differences of hemoglobin values were detected between diabetics and non-diabetics. Here we also demonstrate specific refraction points, which might be able to serve as thresholds for periodontitis.

To summarize, our SoFR-method might add to clinical algorithms in defined study cohorts, since laboratory tests and clinical examination of patients display integral components for medical diagnoses and/or decisions in patients. In particular, data sets from which decisions are deduced for individual patients frequently rely on normal value distributions and depend on a healthy cohort as a reference. While the classical mean ± 1σ method follows the assumption of normal-distribution and thus represents a strict mathematical function, the SoFR-method carries the potential to overcome these limitations in case of non-normal distributed data. Further, cut-off values might be extracted for defined cohorts and diseases, such as periodontitis, and in case of adding another clinical entity to assessment, such as diabetes.

## Conclusion

While the SoFR-method might represent an alternative way to find the threshold for diagnosis, the mean ± k x σ method is well established in diagnostic settings and for definitions of cut-off values deduced from large cohorts of healthy individuals. However, our simple SoFR-method, which can be applied in mixed cohorts with healthy and diseased individuals, might provide good references for suggesting refraction/changing points of the physical status, also for individual patients within a defined race- or disease-defined cohort.

## Author Contributions

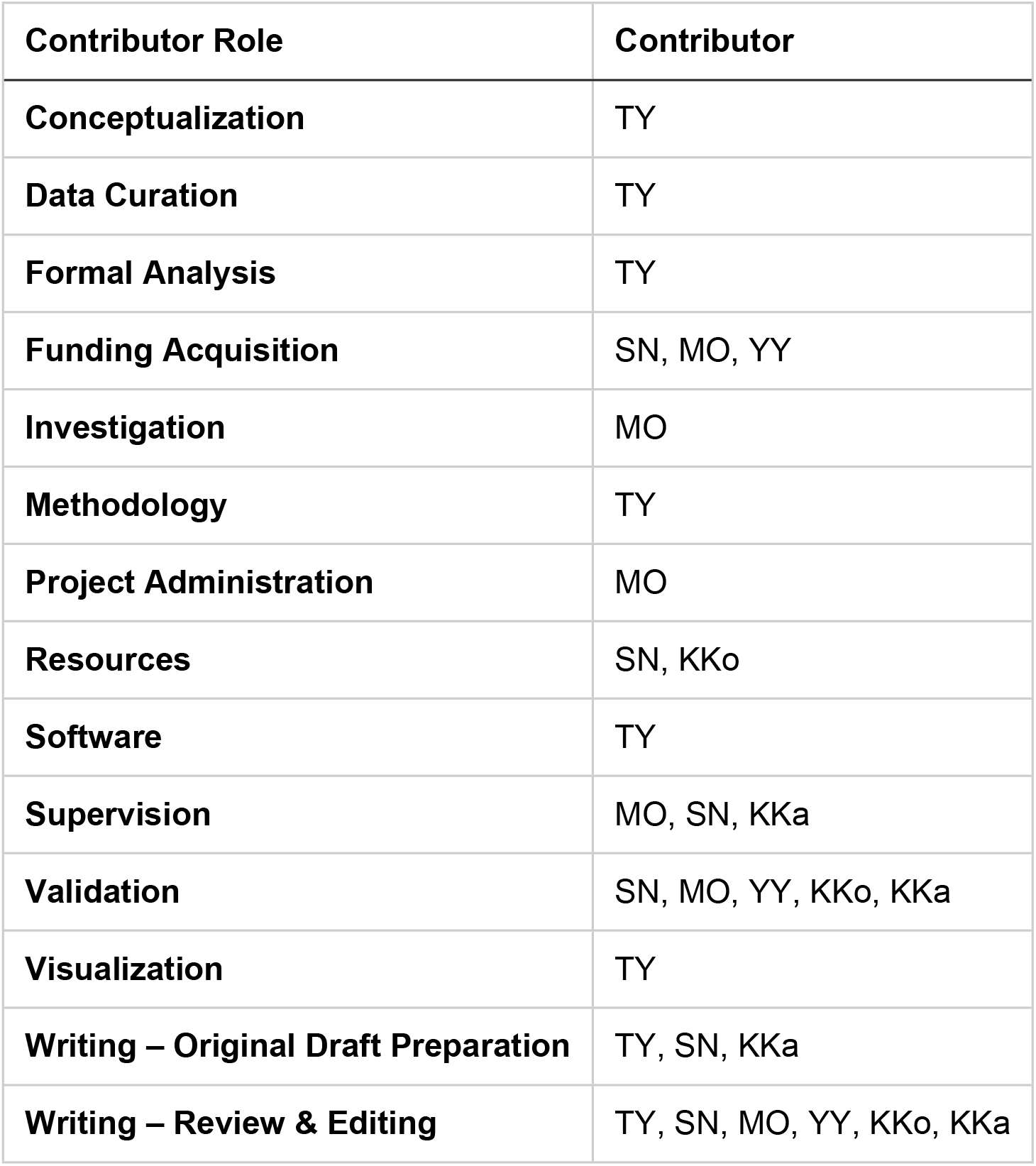

## Acknowledgements

We thank Eiki Matsushita (Hakui General Hospital), Toshihiro Tamura (Tamura Eye Clinic), Tateura Hidemaru (Tateura Dental Clinic), and Naoki Nishida (Naoki Dental Clinic), members of Multi-institutional Collaborative Study in Hakui for their special cooperation. We also thank Masaaki Eto (Ohu University) for his helpful advices.

This study was supported in part by a grant from the Dental Research Center, Nihon University School of Dentistry (2017, 2018), and by a Shika Town Health Promotion Fund, and also supported in part by a grant from Oriental Life Insurance Cultural Development Center.

